# Supramodal executive control of attention: evidence from unimodal and crossmodal dual conflict effects

**DOI:** 10.1101/2020.05.22.110643

**Authors:** Alfredo Spagna, Tingting Wu, Kevin Kim, Jin Fan

**Affiliations:** Department of Psychology, Columbia University in the City University of New York, NY, USA, 10027; Department of Psychology, Queens College, the City University of New York, Queens, NY, USA, 11367

**Keywords:** supramodal executive control, dual conflict, attentional networks, cognitive computational neuroscience

## Abstract

Although we have demonstrated that the executive control of attention acts supramodally as shown by significant correlation between conflict effects measures in visual and auditory tasks, no direct evidence of the equivalence in the computational mechanisms governing the allocation of executive control resources within and across modalities has been found. Here, in two independent groups of 40 participants, we examined the interaction effects of conflict processing in both unimodal (visual) and crossmodal (visual and auditory) dual-conflict paradigms (flanker conflict processing in Task 1 and then in the following Task 2) with a manipulation of the stimulus onset asynchrony (SOA). In both the unimodal and the crossmodal dual-conflict paradigms, the conflict processing of Task 1 interfered with the conflict processing of Task 2 when the SOA was short, reflecting an additive interference effect of Task 1 on Task 2 under the time constraints. These results suggest that there is a unified entity that oversees conflict processing acting supramodally by implementing comparable mechanisms in unimodal and crossmodal scenarios.

## Introduction

In current times, the boundaries of our brain computations are being pushed further and further, with the number of tasks, whether competing or synergistic, steadily increasing. In a hypothetical morning routine, many of us are constantly dealing with conflicting sources of information, all attempting to access our conscious experience. One of the more remarkable functions of our brain is to solve these conflicts, but we often experience a hard limit on how much we can effectively deal with at one time when there is another conflict to resolve. When we find ourselves in a situation that exceeds our computational capacity, interference across concurrent conflict processes may arise, negatively impacting our performance. The reason for such a computational *bottleneck* (Shannon, 1948; Trautwein, Singer, & Kanske, 2016; Wu, Wang, et al., 2019), initially a topic disputed between cognitive psychology and cognitive science (e.g., see Lake, Ullman, Tenenbaum, & Gershman, 2017), is now being investigated under new lenses due to the development of cognitive computational models (Kriegeskorte & Douglas, 2018). Yet, it is still up to debate why humans have a bottleneck, and more importantly, how this computational mechanism becomes responsible for the information-processing limit.

Growing evidence seems to indicate that one of the subcomponents of attention according to a renowned model (Fan, McCandliss, Sommer, Raz, & Posner, 2002; Petersen & Posner, 2012), the executive control, has limited resources and may be a candidate for the above-mentioned computational bottleneck. The executive control of attention is the brain system devoted to inhibit competing messages reaching from sources deemed irrelevant and to support the processing of multisensory inputs from relevant sources (Fan, Flombaum, McCandliss, Thomas, & Posner, 2003; Fan, Fossella, Sommer, Wu, & Posner, 2003; Fan et al., 2009; Fan, Hof, Guise, Fossella, & Posner, 2008; Fan et al., 2007; Martín-Signes, Paz-Alonso, & Chica, 2019; Mullane, Lawrence, Corkum, Klein, & McLaughlin, 2016; Posner, 2012; Spagna, Dong, et al., 2015; Spagna, Kim, Wu, & Fan, 2018; Tian et al., 2016; Trautwein et al., 2016). This system has been shown to act supramodally by integrating stimuli coming from different modalities and cognitive domains, and is in charge of the conflict resolution among competing stimuli to promote an efficient interaction with the environment (Donohue, Liotti, Perez, & Woldorff, 2012; Farah, Wong, Monheit, & Morrow, 1989; Green, Doesburg, Ward, & McDonald, 2011; Ljubojevic et al., 2018; Martín-Signes et al., 2019; Ricciardi, Bonino, Pellegrini, & Pietrini, 2014; Roberts & Hall, 2008; Spagna et al., 2017; Spagna, Mackie, & Fan, 2015). The existence of a unified supramodal function better manages the economic trade-offs inherent in all the biological organisms (Bullmore & Sporns, 2012), while allowing to efficiently coordinate multisensory information as opposed to modality-dependent centres, but comes with the cost of stricter amount of resources available (Kriegeskorte & Douglas, 2018; Spagna, Mackie, et al., 2015).

Yet, the majority of the studies that examined supramodal control functions relied on the examination of the correlation between measurements of the conflict effect as measured by separate visual and auditory conflict effects (e.g., Roberts & Hall, 2008; Spagna et al., 2017; Spagna, Mackie, et al., 2015), providing only indirect evidence for the such claim. For example, we have recently shown that within-subject behavioral measures of the executive control of attention measured by the flanker conflict effect are correlated across a visual and an auditory task (Spagna, Mackie, et al., 2015), and that a deficit of this mechanism is associated with psychiatric disorders that heavily taps on cognitive disfunctions such as schizophrenia (Spagna et al., 2017). Further, conjunction analyses of separate visual and auditory neuroimaging maps relying on the similarities between the spatiotemporal dynamics in the activation of brain areas and networks found during unimodal conflict resolution (e.g., Donohue et al., 2012; Green et al., 2011) provided evidence for the existence of supramodal mechanisms for the executive control. However, the correlational property of our studies as well as of the above-mentioned neuroimaging studies could not provide direct evidence demonstrating that the mechanisms for conflict resolution arising from stimuli in different modalities (i.e., crossmodal conflict) is implemented by the same unit for executive control of attention.

In this study, we tested whether the executive control of attention is the center responsible for handling the interference across competing tasks irrespectively of whether information comes from a single modality or from multiple modalities, i.e., supramodal. We contrasted the behavioral performance in a crossmodal version of the dual conflict paradigm to a unimodal version. The hypothesis that the supramodal executive control of attention is similarly involved in the information processing of both the unimodal and crossmodal conflict resolution would be supported by findings of dual task conflict interference effect in both unimodal and crossmodal paradigms.

## Method

### Participants

A total of 91 undergraduate students of the Psychology Department at Queens College of the City University of New York (CUNY) participated in this study for course requirements and were compensated with course credit. Forty-eight participants completed Experiment 1, while the other 43 participants completed Experiment 2. After excluding those with lower than 70% accuracy, the final sample size was 80, with 40 participants in Experiment 1 (31 females; mean age 20.1 ± 2.9 years; range 18–33 years) and 40 participants in Experiment 2 (26 females; mean age 20.4 ± 3.5 years; range 18–35 years). The protocol was approved by the Institutional Review Board of CUNY, and written informed consent was obtained from each participant prior to participation.

### Task Design

The logic behind the design of the dual conflict paradigm (DCP) is based on the evidence that executing two similar operations simultaneously, e.g., Task 1 and Task 2 (T1 and T2), typically causes interference when participants have to suppress unwanted information (Bourke, 1996, 1997). In the unimodal visual DCP (UV-DCP), the two tasks (T1 and T2) were both visual flanker tasks (Eriksen & Eriksen, 1974; Mackie & Fan, 2016), while in the crossmodal version of the DCP (CM-DCP), the two tasks were a visual flanker task (Fan et al., 2002) and an auditory version of the flanker task also used in a previous study (Spagna, Mackie, et al., 2015). For both tasks, the cross-task interference effect was manipulated by varying the flanker congruency (congruent and incongruent) and interval of presentation of the two tasks (short or long).

#### Experiment 1: Unimodal Visual Dual Conflict Paradigm (UV-DCP)

This task has the same design as in our previous studies (Mackie & Fan, 2016; Spagna et al., 2018). On each trial (**Fig. 1a**), following a 0 to 500 ms randomly varied fixation interval (not shown in Fig. 1 for simplicity), two tasks (T1 and T2) were presented sequentially for 750 ms each with a variable T1 to T2 stimulus onset asynchrony (SOA) of 1000 or 1,00 ms (long and short, respectively). The 750 ms task duration was used to avoid the attentional blink effect, which would interfere with detection of the second target if a shorter (e.g., 500 ms) duration was used. For T1, the stimulus was presented at one of two locations, aligned vertically either above or below the central fixation cross, and consists of a central target arrow flanked by four direction-congruent or incongruent arrows (two on each side) pointing either up or down. For T2, the stimulus was presented either to the left or right of the central fixation cross and includes a central target arrow flanked by four direction-congruent or incongruent arrows, pointing either left or right. T2 was followed by a variable post-target fixation (2,000–2,500 ms). The total time of each trial was 5,000 ms. Participants must sequentially make an up/down response to the central arrows for T1 using the left-hand buttons, and then a left/right response for T2 using the right-hand buttons. For each SOA, four different conflict conditions resulted from the T1-T2 association: CC (T1 congruent, T2 congruent), IC (T1 incongruent, T2 congruent), CI (T1 congruent, T2 incongruent), II (T1 incongruent, T2 incongruent). The task consisted of eight blocks in total, with 64 trials per block. Each block lasted approximately 6 min, and the total task time was approximately 50 min. Participants were instructed to respond as quickly and accurately as possible and take self-initiated and self-terminated breaks as needed between blocks to control for fatigue.

A previous study using the UV-DCP (Mackie & Fan, 2016) demonstrated that under the 1,000 ms SOA, the computational load of T2 was not significantly affected by T1, indicating that the information processing involved in each task performs sequentially without an overlap. However, under the 100 ms SOA, the tasks occur in much quicker succession, resulting in task processing overlap, with the computational load during T2 period approaching the sum of the computational load of T1 and T2, an additive effect based on RT pattern.

**Figure 1.**
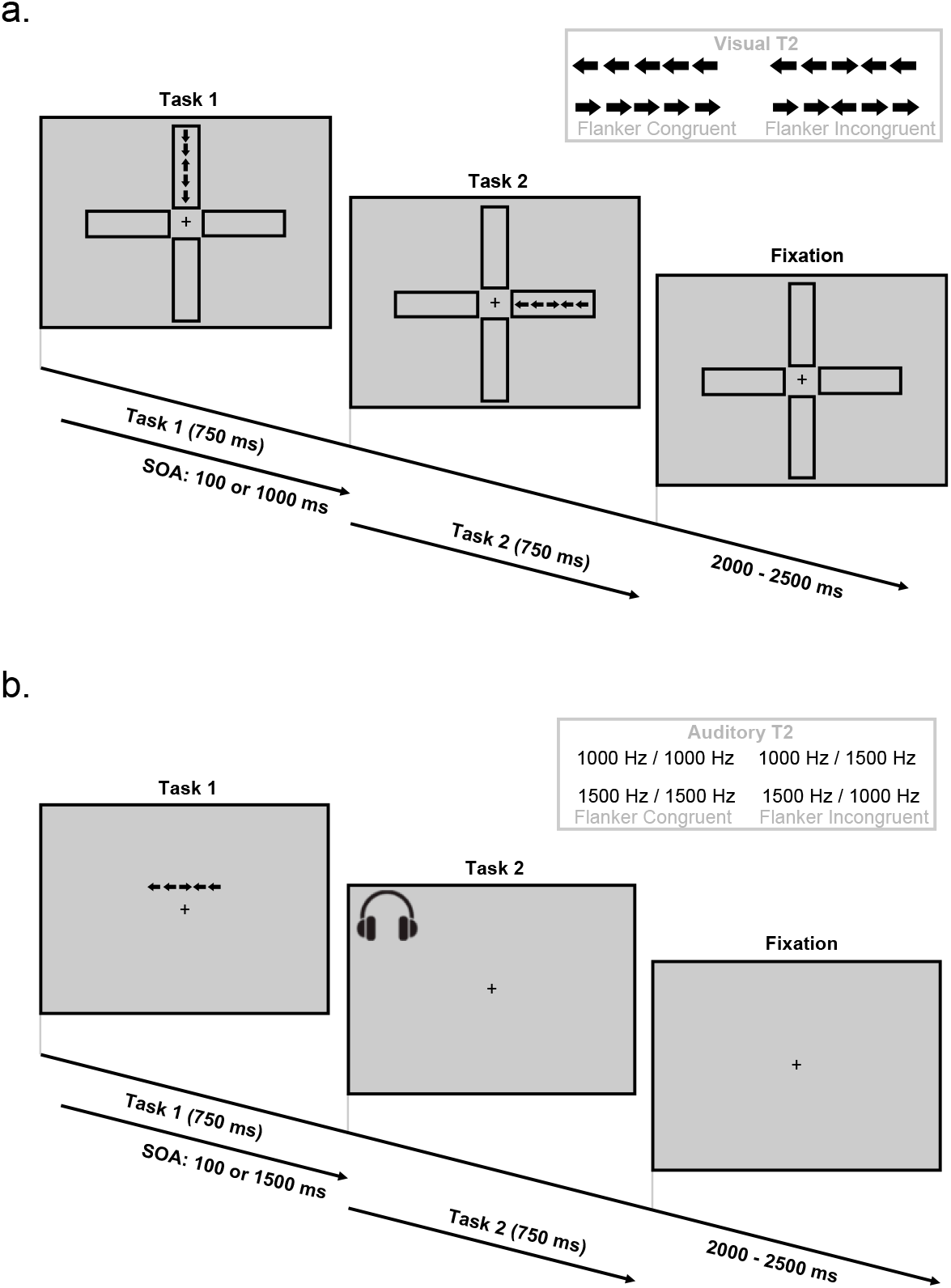
(a) Schematic of the UV-DCP. Participants were presented with two flanker tasks in succession and were required to indicate the direction of the center arrow and to ignore the flanker arrows, which may either be congruent or incongruent with the target. Stimulus onset asynchrony was manipulated such that Task 2 appeared either 100 or 1,000 ms after Task 1. (b) Schematic of the CM-DCP. Participants were presented with two flanker tasks in succession. For the visual task, participants were required to indicate the direction of the center arrow and ignore the flanker arrows; for the auditory task, participants were required to indicate the duration of the first tone (the target) and ignore the second (distracting tone) in the auditory task. The distracting tone was either congruent or incongruent with the target. Stimulus onset asynchrony was manipulated such that Task 2 appears either 100 or 1,500 ms after Task 1.

#### Experiment 2: Crossmodal Dual Conflict Task (CM-DCP)

On each trial (**Fig. 1b**), following a 0 to 500 ms randomly varied fixation interval, T1 and T2 were presented sequentially for 750 ms each with a variable T1 to T2 SOA of 100 or 1500 ms. As in the UV-DCP, under the 100 ms SOA, the tasks occur in much quicker succession, resulting in task processing overlap, with the computational load during T2 processing approaching the sum of the computational load of T1 and T2, as an additive effect based on RT pattern. There were two block orders: in the V-A blocks, the visual task was presented first, followed by the auditory task, while in the A-V blocks, the order was reversed. For the visual task, as in Experiment 1, the target was an arrow presented one degree above or below the central fixation point flanked by four direction-congruent or incongruent arrows (two on each side) pointing either left or right. For the auditory task (low/high pitch judgment), the target was a binaurally presented tone with a 100 ms duration followed by a 100 ms delay then a second pitch-congruent or incongruent tone presented for 100 ms, with tones being either 1000 or 1500 Hz. Participants were also instructed to maintain the order of task presentation by responding to T1 first and then T2 sequentially, and to make a left or right response according to the direction of the central arrows for the visual task using the right hand buttons and an up/down response according to the high or low pitch of the target for the auditory task using the left hand buttons. In parallel with Experiment 1, for each SOA, there were four different conflict conditions resulted from the T1-T2 association: CC (T1 congruent, T2 congruent), IC (T1 incongruent, T2 congruent), CI (T1 congruent, T2 incongruent), II (T1 incongruent, T2 incongruent). T2 was followed by a variable post-target fixation (2,000-2,500 ms), with total trial time of 4,300 ms. There were eight blocks, four V-A and four A-V, each composed of 64 trials. Each block lasted approximately 9 mins, and the total task time was approximately 70 mins. Participants were instructed to respond as quickly and accurately as possible and take self-initiated and self-terminated breaks as needed between runs to control for fatigue.

### Data Analysis

#### Analysis of Variance (ANOVA) on T1- and T2-locked Reaction Time and Error Rate

Data from each dual conflict paradigm were analyzed separately using ANOVA, with task conditions as the within subject factors. Data were first tested for normal distributions, homogeneity of variance, and assumption of sphericity. None of the above-mentioned assumptions was violated. For each experiment, T1-locked and T2-locked RT and error rate scores underwent a 2 (*SOA*: long, short) × (*T1 Congruency*: Congruent, Incongruent) × 2 (*T2 Congruency*: Congruent, Incongruent) ANOVA to examine the dual conflict effects in the visual and crossmodal conditions. The main effect of the factor SOA was examined by comparing participants’ performance in trials with long *vs*. short SOA. The main effect of the factor T1 Congruence was examined by comparing participant’s performance in trials with T1 incongruent *vs* T1 congruent. The main effect of the factor T2 Congruence was examined by comparing participant’s performance in trials with T2 incongruent *vs* T2 congruent. The SOA × T1 interaction effect was especially examined for the interference of T1 congruency on T2, with a significant interaction indicating that the computational load of T1 has greater disruptive effect on T2-lock performance between two SOAs. Specifically, if the processing of T1 overlaps with the processing of T2, even if the two tasks were from different modalities, the computational load during T1 carries over the processing of T2, approaching the sum of the computational load of Tasks 1 and 2, producing an additive effect based on T2-locked RT pattern. This effect, if significant in both unimodal and crossmodal tasks, would provide the evidence of supramodal executive control. However, if the competition between T1 and T2 tasks requires additional resources when the tasks are performed concurrently, hindering participants’ performance to a greater extent, there should be a three-way interaction between the cognitive loads in the T1 and T2 tasks at short SOA and when both T1 and T2 conditions were incongruent. A critical alpha level of .05 was used.

### Additive effect of cognitive load across tasks

The information theory account of cognitive control (Fan, 2014; Mackie & Fan, 2016; Wu et al., 2017; Wu, Wang, et al., 2019) allowed us to estimate the computational load for a binary response as approximately 1 bit (log_2_(2) for the two possible response alternatives in each flanker task) under the congruent condition, and greater than 1 but less than 2 bits under the incongruent condition. The difference between incongruent and congruent conditions in terms of bits would, therefore, be less than or approximately 1. In addition, on each trial the target stimulus could be presented in two alternative locations (for the visual task) or in two alternative frequency ranges (for the auditory task). Such variation contributes to an additional computational load of 1 bit. Therefore, when T1 and T2 are sufficiently separated in time (long SOA), they can be performed sequentially, and the resulting minimum and maximum computational loads for both tasks would be 2 (congruent condition) and 3 (incongruent condition) bits, respectively.

## Results

**Figure 2**. shows T1-locked and T2-locked performance separately for the UV-DCP (left two columns) and CM-DCP (right two columns) at each level of the SOA.

**Figure 2.**
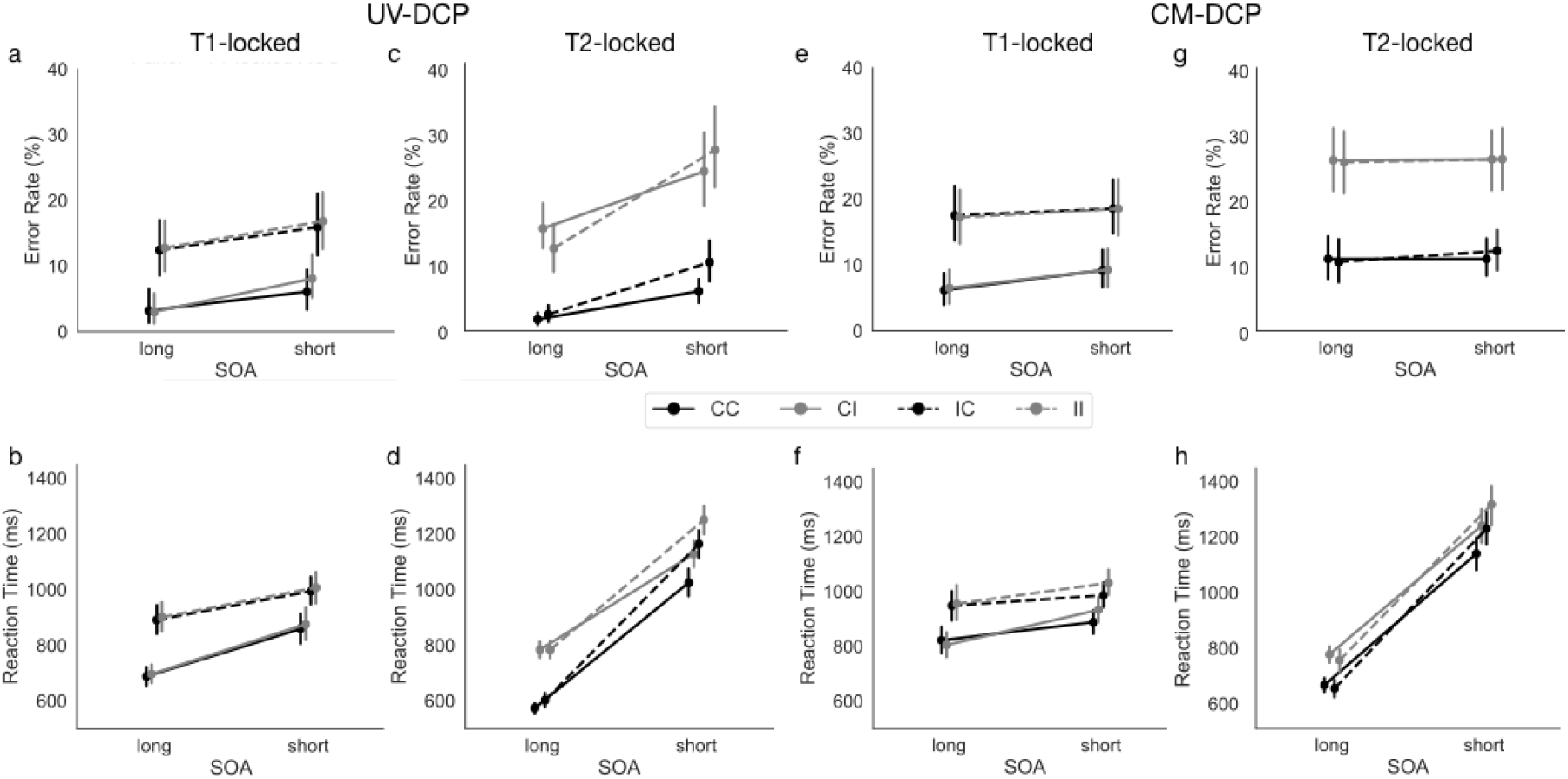
Results from the ANOVA conducted on T1-locked a) Error rate (in percentage) and b) Reaction Time (in ms) and T2-locked c) Error rate (in percentage) and d) Reaction Time (in ms) for the UV-DCP task; e) Error rate (in percentage) and f) Reaction Time (in ms) and T2-locked g) Error rate (in percentage) and h) Reaction Time (in ms) for the CM-DCP task.

### The dual conflict effects in the visual modality

#### Results on T1-locked performance

**Figure 2a** and **2b** show a graphical representation of the T1-locked effects. For error rate of T1, the main effect of the factor *SOA* was significant (*F*_(1,39)_ = 22.61, *p* < .001, *η*_*2*_ = .37), with more errors in the short SOA condition (11.74 ± 12.97) compared to the long SOA condition (7.83 ± 11.90%). The main effect of the factor *T1 Congruency* was significant (*F*_(1,39)_ = 45.68, *p* < .001, *η*_*2*_ = .54), with more errors in the incongruent condition (14.49 ± 14.00%) compared to the congruent condition (5.08 ± 9.40%). The main effect of the factor *T2 Congruency* was not significant (*F*_(1,39)_ = 3.21, *p* = .08), while the interaction between *SOA* and *T2 Congruency* was significant (*F*_(1,39)_ = 4.65, *p* < .05, *η*_*2*_ = .11). This interaction showed that T2 conflict effect was greater at short SOA (T2 congruent: 11.02 ± 14.13% *vs*. T2 incongruent condition 12.46 ± 12.69%, *p* < .05, compared to the long SOA condition, *p* = .87), indicating that the presentation of T2-task conditions interfered with the processing of T1. The interaction between *SOA* and *T1 Congruency* (F <1), between *T1 Congruency* and *T2 Congruency* (F <1), and the three-way interaction (*F*_(1,39)_ = 1.77, *p* = .19) were not significant.

For RT of T1, the analysis showed that the main effect of the factor *SOA* was significant (*F*_(1,39)_ = 85.25, *p* < .001, *η*_*2*_ = .69), with slower RTs in the short SOA condition (934 ± 201ms) compared to the long SOA condition (794 ± 175ms). The main effect of the factor *T1 Congruency* was significant (*F*_(1,39)_ = 169.94, *p* < .001, *η*_*2*_ = .81), with slower RTs in the incongruent condition (949 ± 179ms) compared to the incongruent condition (780 ± 172ms). The main effect of the factor *T2 Congruency* was significant (*F*_(1,39)_ = 15.76, *p* < .001, *η*_*2*_ = .29), with slower RT in the incongruent condition (870 ± 197ms) compared to the congruent condition (858 ± 199ms). The interaction between *SOA* and *T1 Congruency* was also significant (*F*_(1,39)_ = 23.31, *p* < .001, *η*_*2*_ = .37), with smaller T1 conflict effect short SOA (T1 congruent: 867 ± 180 ms *vs*. T1 incongruent: 1001 ± 174 ms, *p* < .05), compared to the conflict effect at long SOA condition (T1 congruent: 692 ± 108ms *vs* T1 incongruent: 896 ± 170ms, *p* <.01), indicating that the processing of T1 conflict was more efficient for the short SOA. The interaction between *SOA* and *T2 Congruency* (F <1), between *T1 Congruency* and *T2 Congruency* (F <1), and the three-way interaction (F< 1) were not significant.

#### Results on T2-locked performance

**Figure 2c** and **2d** show a graphical representation of the T2-locked results. For error rate of T2, the main effect of the factor *SOA* was significant (*F*_(1,39)_ = 22.61, *p* <.001, *η*_*2*_ = .47), with more errors in the short SOA condition (16.74 ± 16.52%) compared to the long SOA condition (7.91 ± 10.42%). The main effect of the factor *T1 Congruency* was not significant (*F*_(1,39)_ = 3.10, *p* = .09). The main effect of the factor *T2 Congruency* was significant (*F*_(1,39)_ = 79.88, *p* < .001, *η*_*2*_ = .67), with more errors in the incongruent condition (19.60 ± 7.51%) compared to the congruent condition (5.05 ± 16.83%). The interaction between *SOA* and *T1 Congruency* was also significant (*F*_(1,39)_ = 17.79, *p* < .001, *η*_*2*_ = .31), demonstrating an additive effect, with T1 incongruent condition having a greater impact on error rate at short SOA (T1 congruent: 14.86 ± 16.39% *vs* T1 incongruent: 18.61 ± 17.75%, *p* < .05) compared to long SOA (T1 congruent: 8.46 ± 7.24 *vs* T1 incongruent: 7.46 ± 7.98, *p* = .08). The interaction between *SOA* and *T2 Congruency* was also significant (*F*_(1,39)_ = 7.84, *p* < .01, *η*_*2*_ = .17), with the conflict effect in terms of error rate greater at short SOA (T2 congruent: 8.07 ± 8.47% *vs*. T2 incongruent: 25.41 ± 19.15%, *p* < .05) compared to the long SOA (T2 congruent: 2.03 ± 3.67% *vs* T2 incongruent: 13.07 ± 11.62%, *p* < .01). The interaction between *T1 Congruency* and *T2 Congruency* was also significant (*F*_(1,39)_ = 6.73, *p* < .05, *η*_*2*_ = .15), with a greater T1 conflict effect at T2 congruent (T1 congruent: 3.77 ± 4.53 % *vs*. T1 incongruent: 6.33 ± 10.71%, *p* < .05) compared to T2 incongruent condition (T1 congruent: 19.55 ± 14.98% *vs*. T1 incongruent: 19.65 ± 15.80%, *p* < .05). The three-way interaction was not significant (*F*_(1,39)_ = 3.25, *p* = .08).

For RT of T2, the main effect of the factor *SOA* was significant (*F*_(1,39)_ = 941.15, *p* < .001, *η*_*2*_ = .96), with slower RT in the short SOA condition (1143 ± 281 ms) compared to the long SOA condition (686 ± 131 ms). The main effect of the factor *T1 Congruency* was significant (*F*_(1,39)_ = 101.65, *p* < .001, *η*_*2*_ = .72), with slower RT in the incongruent condition (951 ± 300 ms) compared to the congruent condition (878 ± 250 ms). The main effect of the factor *T2 Congruency* was significant (*F*_(1,39)_ = 434.28, *p* < .001, *η*_*2*_ = .92), with slower RT in the incongruent condition (987 ± 295 ms) compared to the incongruent condition (841 ± 247 ms). The interaction between *SOA* and *T1 Congruency*, which was the focus of this study, was also significant (*F*_(1,39)_ = 111.62, *p* < .001, *η*_*2*_ = .74). In line with results on T2-locked error rate, this interaction demonstrated an additive effect, with T1 incongruent condition having a greater impact on RT of T2 at short SOA (T1 congruent: 1077 ± 167 ms *vs*. T1 incongruent: 1209 ± 171 ms, *p* < .01) compared to long SOA (T1 congruent 679 ± 132 ms *vs*. T1 incongruent: 693 ± 130 ms, *p* < .05). The interaction between *SOA* and *T2 Congruency* was also significant (*F*_(1,39)_ = 139.17, *p* < .001, *η*_*2*_ = .78). Diverging from the results on T2-locked error rate for this interaction, the conflict effect in RT was less at short SOA (T2 congruent: 1095 ± 179 ms *vs*. T2 incongruent: 1191 ± 170 ms, *p* < .01) compared to long SOA (T2 congruent: 587 ± 74 ms *vs*. T2 incongruent: 783 ± 98 ms, *p* < .01), indicating the presence of a speed-accuracy trade-off. The interaction between *T1 Congruency* and *T2 Congruency* was also significant (*F*_(1,39)_ = 10.72, *p* < .001, *η*_*2*_ = .22). In line with results on T2-locked error rate, this interaction showed that the T1 conflict effect was less at T2 incongruent (T1 congruent: 956 ± 123 ms *vs*. T1 incongruent: 1017 ± 135 ms, *p* < .01) compared to T2 congruent (T1 congruent: 799 ± 112 ms *vs*. T1 incongruent: 883 ± 124.5 ms, *p* < .01). The three-way interaction was not significant (*F* < 1).

### The dual conflict effects across the visual and auditory modalities

#### Results on T1-locked performance

**Figure 2 e** and **f** show a graphical representation of the results of T1 in error rate and RT. For error rate of T1, the main effect of the factor *SOA* was significant (*F*_(1,39)_ = 23.60, *p* <.001, *η*_*2*_ = .38), with more errors in the short SOA condition (13.91 ± 12.20%) compared to the long SOA condition (11.88 ± 12.55%). The main effect of the factor *T1 Congruency* was significant (*F*_(1,39)_ = 38.45, *p* <.001, *η*_*2*_ = .50), with more errors in the incongruent condition (17.99 ± 13.60%) compared to the congruent condition (7.8 ± 8.95%). The main effect of the factor *T2 Congruency* was not significant (F_(1,39)_ = 3.21, *p* = .08). The interaction between *SOA* and *T1 Congruency* was significant (F_(1,39)_ = 6.18, *p* <.05, *η*_*2*_ = .14), with the T1 conflict effect smaller at short SOA (T1 congruent: 9.25 ± 9.42% *vs* T1 incongruent: 18.57 ± 13.62%, *p* <.05), while this difference was only marginally significant for the long SOA condition (T1 congruent: 6.35 ± 8.32% *vs* T1 incongruent: 17.41 ± 13.73%, *p* = .05). The interaction between *SOA* and *T2 Congruency* (F < 1), between *T1 Congruency* and *T2 Congruency* (F < 1), and the three-way interaction (F < 1) were not significant.

For RT of T1, the main effect of the factor *SOA* was significant (*F*_(1,39)_ = 31.22, *p* <.001, *η*_*2*_ = .44), with slower RT in the short SOA condition (960 ± 166ms) compared to the long SOA condition (883 ± 185ms). The main effect of the factor *T1 Congruency* was significant (*F*_(1,39)_ = 224.93, *p* <.001, *η*_*2*_ = .85), with slower RT in the incongruent condition (980 ± 176ms) compared to the congruent condition (862 ± 155ms). The main effect of the factor *T2 Congruency* was significant (*F*_(1,39)_ = 7.30, *p* < .01, *η*_*2*_ = .16), with slower RT in the incongruent condition (931 ± 174ms) compared to the congruent condition (911 ± 185ms). In line with results from the T1-locked error rate, the interaction between *SOA* and *T1 Congruency* was also significant (*F*_(1,39)_ = 15.75, *p* < .001, *η*_*2*_ = .29). This interaction showed that the T1 conflict effect was smaller at short SOA (T1 congruent: 911 ± 145ms *vs* T1 incongruent: 1009 ± 154ms, *p* <.01), compared to the long SOA (T1 congruent: 814 ± 150ms *vs* T1 incongruent: 952 ± 192ms, *p* <.01). The interaction between *SOA* and *T2 Congruency* was also significant (*F*_(1,39)_ = 47.06, *p* < .001, *η*_*2*_ = .55). This interaction showed that T2 conflict effect was greater at short SOA (T2 congruent: 937 ± 150ms *vs* T2 incongruent: 982 ± 162ms, *p* <.01, compared to long SOA (*p* = .57), indicating that the presentation of T2-task conditions interfered with the processing of T1. The interaction between *T1 Congruency* and *T2 Congruency* (*F*_(1,39)_ = 2.16, *p* = .15), and the three-way interaction (*F*_(1,39)_ = 2.43, *p* = .13) were not significant.

#### Results on T2-locked performance

**Figure 2g** and **2h** show a graphical representation of T2-locked results in error rate and RT. For error rates of T2, the main effect of the factor *SOA* (*F*_(1,39)_ = 1.05, *p* = .32), and *T1 Congruency* (*F* < 1) were not significant. The main effect of the factor *T2 Congruency* was significant (*F*_(1,39)_ = 88.21, *p* < .001, *η*_*2*_ = .69), with more errors in the incongruent condition (25.95 ± 10.72%) compared to the congruent condition (11.12 ± 14.99%). The interaction between *SOA* and *T1 Congruency* (*F*_(1,39)_ = 2.5, *p* = .12), SOA and *T2 Congruency* (F <1), between *T1 Congruency* and *T2 Congruency* (*F*_(1,39)_ = 1.35, *p* = .25), and the three-way interaction (*F*_(1,39)_ =1.4, *p* = .24) were not significant.

For RT of T2, the analysis showed that the main effect of the factor *SOA* was significant (*F*_(1,39)_ = 498.39, *p* <.001, *η*_*2*_ = .93), with slower RT in the short SOA condition (1230 ± 301 ms) compared to the long SOA condition (705 ± 119 ms). The main effect of the factor *T1 Congruency* was significant (*F*_(1,39)_ = 28.95, *p* <.001, *η*_*2*_ = .43), with slower RT in the incongruent condition (984 ± 338 ms) compared to the congruent condition (950 ± 290 ms). The main effect of the factor *T2 Congruency* was significant (*F*_(1,39)_ = 254.72, *p* < .001, *η*_*2*_ = .87), with slower RT in the incongruent condition (1018 ± 306 ms) compared to the congruent condition (917 ± 312 ms). Most importantly, the interaction between *SOA* and *T1 Congruency* was significant (*F*_(1,39)_ = 111.62, *p* < .001, *η*_*2*_ = .74) as an additive effect, with T1 incongruent condition having a greater impact on RT at short SOA (T1 congruent: 1188 ± 210 ms *vs*. T1 incongruent: 1272 ± 211 ms, *p* < .01) compared to long SOA (T1 congruent: 713 ± 91 ms *vs*. T1 incongruent: 696 ± 119 ms, *p* = .07). The interaction between *SOA* and *T2 Congruency* (*F*_(1,39)_ = 1.23, *p* = .27), between *T1 Congruency* and *T2 Congruency* (*F*_(1,39)_ = 3.71, *p* = .06), and the three-way interaction (*F <1*) were not significant.

## Discussion

Moving beyond previous correlational evidence of conflict resolution measures in unimodal tasks (e.g., Spagna et al., 2017; Spagna, Mackie, et al., 2015), this study provides direct evidence that supramodal function of the executive control of attention supports dual-task conflict performance by showing that the interference produced by the dual conflict paradigm follows similar trajectories in unimodal and crossmodal paradigms.

Results from this study support our hypothesis regarding the existence of a resource-limited central capacity unit in the brain (Tombu et al., 2011; Tombu & Jolicœur, 2003) dealing constantly with information coming from different sensory modalities. Here, we further expand on this central capacity model, by proposing that the interference effect results from the necessity of a single unified supramodal center.

Compared to multiple modality-specific centers, this unit best solves the economic trade-off between the benefits of adaptively and efficiently coordinating information from multiple modalities and the costs of limited capacity (Dehaene, Lau, & Kouider, 2017; Fan, 2014; Mackie & Fan, 2016; Wu et al., 2017; Wu, Wang, et al., 2019). Furthermore, the additive effect found in our results suggests that a single center is necessary and sufficient to efficiently coordinate information from different modalities.

The behavioral pattern found in this study uncovers the mental algorithm implemented while processing two concurrent conflict tasks (i.e., when T1 overlaps with the processing of T2), as shown by the additive interaction effect that approaches the sum of the computational loads of each task. This pattern manifests as a significant decrease in performance observed in T2-locked reaction time and error rates, indicating the computational load during T1 carries over the processing of T2, and that the limited resources can be distributed only serially to solve the first conflict at hand. Importantly for the scope of this present study, we demonstrated that the same additive pattern can be identified irrespectively of the sensory modality of the stimuli. This result is in line with long-standing literature (Shannon, 1948; Trautwein et al., 2016; Wu, Wang, et al., 2019) suggesting the existence of a computational bottleneck limiting the amount of information that can be processed at any given moment, and seems to unequivocally indicate that this bottleneck is in the function of the supramodal executive control of attention (Spagna et al., 2017; Spagna, Mackie, et al., 2015). Specifically, the need to resolve the conflict among competing stimuli and promoting an efficient interaction with the environment triggers the function of the supramodal executive control of attention, but the limited resources inherent to the presence of a single unit in the brain that manages multisensory information comes at a cost of decreased performance when resources are exhausted.

The role of the executive control of attention as a supramodal unit is also supported by existing neuroimaging literature. For instance, long standing evidence has shown that activity in brain regions spanning across the fronto-parietal (Fan et al., 2014) and cingulo-opercular (Dosenbach, Fair, Cohen, Schlaggar, & Petersen, 2008) networks is associated with the resolution of conflict among competing stimuli, irrespectively of whether they are presented in the visual (Fan, McCandliss, Fossella, Flombaum, & Posner, 2005; Petersen & Posner, 2012; Xuan et al., 2016) or auditory modality (Donohue et al., 2012; Farah et al., 1989; Green et al., 2011; Roberts & Hall, 2008). The critical role of this system in promoting an efficient interaction with the environment is also supported by recent meta-analytic evidence showing the involvement of these same fronto-parietal and cingulo-opercular networks in various control functions spanning across the domain of cognitive control, working memory, and decision making (Wu, Chen, et al., 2019). Taken together, this wealth of knowledge might support an even greater expansion of the model proposed, indicating that to process multisensory inputs from relevant sources and to inhibit competing messages deemed irrelevant, a supramodal and *supra-domain* system exerting top-down control is necessary, which we propose is the executive control of attention.

Investigating the computational mechanism governing the supramodal executive control of attention can benefit our understanding of many neurological and psychiatric disorders. Deficits in the integration of multisensory information and decreases in dual task performance have been shown in patients with autism spectrum disorder (Mackie & Fan, 2016), patients with unilateral neglect (Bolognini et al., 2016), patients with Parkinson Disease (Strouwen et al., 2015), and patients with schizophrenia (see also Helfrich & Knight, 2019; Spagna et al., 2017). It is highly likely that the aberrant supramodal mechanism found in these neurological or psychiatric conditions actually masks different patterns of inefficient coordination of information flow from sensory areas to association cortex. For instance, in our recent study (Spagna et al., 2017) we showed that while healthy controls show a significant correlation between conflict effects in the visual and auditory modality, an indication of an intact supramodal executive control mechanism, patients with schizophrenia do not show this correlation. This deficit was associated with the patients’ clinical symptoms, and that patients’ performance was mostly impaired when both the supramodal and the modality-specific (i.e., visual or auditory) mechanisms were impaired. The intactness of the supramodal mechanism of the executive control of attention depends on the normal functioning of brain attention networks, and that either lesion (e.g., in the case of the neglect) or hypo-activation of nodes in these brain network (as shown in schizophrenia) would result in a peculiar clinical symptomatology based on the specific disconnection pattern at hand.

In conclusion, the amount of evidence that our group, as well as others (e.g., see Helfrich & Knight, 2019 for a review) have collected over the years, seems to unequivocally favor the notion that the executive control function is the attentional component that allows individuals to solve conflict arising from various modalities, by integrating priors, incoming information, and contextual rules in the service of goal-directed behavior.

## Author Contributions

J.F., A.S., and T.W. designed the experiments; A.S. analyzed the data. All authors discussed the results and contributed to writing up the report. Behavioral data and the code to reproduce the figures (built using Python on Spyder) can be found on the GitHub page of A.S.

## Reference List

Bolognini, N., Convento, S., Casati, C., Mancini, F., Brighina, F., & Vallar, G. (2016). Multisensory integration in hemianopia and unilateral spatial neglect: Evidence from the sound induced flash illusion. Neuropsychologia, 87, 134–143.

Bourke, P. A. (1996). A general factor involved in dual task performance decrement. The Quarterly Journal of Experimental Psychology: Section A, 49(3), 525–545.

Bourke, P. A. (1997). Measuring attentional demand in continuous dual-task performance. The Quarterly Journal of Experimental Psychology Section A, 50(4), 821–840.

Bullmore, E., & Sporns, O. (2012). The economy of brain network organization. Nat Rev Neurosci, 13(5), 336–349. doi: 10.1038/nrn3214

Dehaene, S., Lau, H., & Kouider, S. (2017). What is consciousness, and could machines have it? Science, 358(6362), 486–492.

Donohue, S. E., Liotti, M., Perez, R., 3rd, & Woldorff, M. G. (2012). Is conflict monitoring supramodal? Spatiotemporal dynamics of cognitive control processes in an auditory Stroop task. Cogn Affect Behav Neurosci, 12(1), 1–15. doi: 10.3758/s13415-011-0060-z

Dosenbach, Fair D. A., Cohen, A. L., Schlaggar, B. L., & Petersen, S. E. (2008). A dual-networks architecture of top-down control. Trends Cogn Sci, 12(3), 99–105. doi: 10.1016/j.tics.2008.01.001

Eriksen, B. A., & Eriksen, C. W. (1974). Effects of noise letters upon the identification of a target letter in a nonsearch task. Perception & psychophysics, 16(1), 143–149.

Fan, J. (2014). An information theory account of cognitive control. Frontiers in human neuroscience, 8, 680–680.

Fan, J., Flombaum, J. I., McCandliss, B. D., Thomas, K. M., & Posner, M. I. (2003). Cognitive and brain consequences of conflict. NeuroImage, 18(1), 42–57. doi: 10.1006/nimg.2002.1319

Fan, J., Fossella, J., Sommer, T., Wu, Y., & Posner, M. I. (2003). Mapping the genetic variation of executive attention onto brain activity. Proc Natl Acad Sci U S A, 100(12), 7406–7411. doi: 10.1073/pnas.0732088100

Fan, J., Gu, X., Guise, K. G., Liu, X., Fossella, J., Wang, H., & Posner, M. I. (2009). Testing the behavioral interaction and integration of attentional networks. Brain and Cognition, 70(2), 209–220. doi: 10.1016/j.bandc.2009.02.002

Fan, J., Hof, P. R., Guise, K. G., Fossella, J. A., & Posner, M. I. (2008). The functional integration of the anterior cingulate cortex during conflict processing. Cereb Cortex, 18(4), 796–805. doi: 10.1093/cercor/bhm125

Fan, J., Kolster, R., Ghajar, J., Suh, M., Knight, R. T., Sarkar, R., & McCandliss, B. D. (2007). Response anticipation and response conflict: an event-related potential and functional magnetic resonance imaging study. J Neurosci, 27(9), 2272–2282. doi: 10.1523/JNEUROSCI.3470-06.2007

Fan, J., McCandliss, B. D., Fossella, J., Flombaum, J. I., & Posner, M. I. (2005). The activation of attentional networks. Neuroimage, 26(2), 471–479. doi: 10.1016/j.neuroimage.2005.02.004

Fan, J., McCandliss, B. D., Sommer, T., Raz, A., & Posner, M. I. (2002). Testing the efficiency and independence of attentional networks. J Cogn Neurosci, 14(3), 340–347. doi: 10.1162/089892902317361886

Fan, J., Van Dam, N. T., Gu, X., Liu, X., Wang, H., Tang, C. Y., & Hof, P. R. (2014). Quantitative characterization of functional anatomical contributions to cognitive control under uncertainty. Journal of Cognitive Neuroscience, 26(7), 1490–1506. doi: 10.1162/jocn_a_00554

Farah, M. J., Wong, A. B., Monheit, M. A., & Morrow, L. A. (1989). Parietal lobe mechanisms of spatial attention: Modality-specific or supramodal? Neuropsychologia, 27(4), 461–470.

Green, J. J., Doesburg, S. M., Ward, L. M., & McDonald, J. J. (2011). Electrical neuroimaging of voluntary audiospatial attention: evidence for a supramodal attention control network. J Neurosci, 31(10), 3560–3564. doi: 10.1523/JNEUROSCI.5758-10.2011

Helfrich, R. F., & Knight, R. T. (2019). Cognitive neurophysiology of the prefrontal cortex Handbook of clinical neurology (Vol. 163, pp. 35–59): Elsevier.

Kriegeskorte, N., & Douglas, P. K. (2018). Cognitive computational neuroscience. Nature neuroscience, 21(9), 1148–1160.

Lake, B. M., Ullman, T. D., Tenenbaum, J. B., & Gershman, S. J. (2017). Building machines that learn and think like people. Behavioral and brain sciences, 40.

Ljubojevic, V., Luu, P., Gill, P. R., Beckett, L.-A., Takehara-Nishiuchi, K., & De Rosa, E. (2018). Cholinergic modulation of frontoparietal cortical network dynamics supporting supramodal attention. Journal of Neuroscience, 38(16), 3988–4005.

Mackie, M. A., & Fan, J. (2016). Reduced Efficiency and Capacity of Cognitive Control in Autism Spectrum Disorder. Autism Research, 9(3), 403–414. doi: 10.1002/aur.1517

Martín-Signes, M., Paz-Alonso, P. M., & Chica, A. B. (2019). Connectivity of frontoparietal regions reveals executive attention and consciousness interactions. Cerebral Cortex, 29(11), 4539–4550.

Mullane, J. C., Lawrence, M. A., Corkum, P. V., Klein, R. M., & McLaughlin, E. N. (2016). The development of and interaction among alerting, orienting, and executive attention in children. Child Neuropsychol, 22(2), 155–176. doi: 10.1080/09297049.2014.981252

Petersen, S. E., & Posner, M. I. (2012). The attention system of the human brain: 20 years after. Annu Rev Neurosci, 35, 73–89. doi: 10.1146/annurev-neuro-062111-150525

Posner, M. I. (2012). Imaging attention networks. Neuroimage, 61(2), 450–456. doi: 10.1016/j.neuroimage.2011.12.040

Ricciardi, E., Bonino, D., Pellegrini, S., & Pietrini, P. (2014). Mind the blind brain to understand the sighted one! Is there a supramodal cortical functional architecture? Neuroscience & Biobehavioral Reviews, 41, 64–77.

Roberts, K. L., & Hall, D. A. (2008). Examining a supramodal network for conflict processing: a systematic review and novel functional magnetic resonance imaging data for related visual and auditory stroop tasks. Journal of cognitive neuroscience, 20(6), 1063–1078.

Shannon, C. E. (1948). A mathematical theory of communication. Bell system technical journal, 27(3), 379–423.

Spagna, A., Dong, Y., Mackie, M.-A., Li, M., Harvey, P. D., Tian, Y., … Fan, J. (2015). Clozapine improves the orienting of attention in schizophrenia. Schizophrenia Research, 168(1), 285–291. doi: 10.1016/j.schres.2015.08.009

Spagna, A., He, G., Jin, S., Gao, L., Mackie, M. A., Tian, Y., … Fan, J. (2017). Deficit of supramodal executive control of attention in schizophrenia. J Psychiatr Res, 97, 22–29. doi: 10.1016/j.jpsychires.2017.11.002

Spagna, A., Kim, T. H., Wu, T., & Fan, J. (2018). Right hemisphere superiority for executive control of attention. Cortex. doi: 10.1016/j.cortex.2018.12.012

Spagna, A., Mackie, M.-A., & Fan, J. (2015). Supramodal Executive Control of Attention. Frontiers in Psychology, 6. doi: 10.3389/fpsyg.2015.00065

Strouwen, C., Molenaar, E. A., Münks, L., Keus, S. H., Bloem, B. R., Rochester, L., & Nieuwboer, A. (2015). Dual tasking in Parkinson’s disease: should we train hazardous behavior? Expert review of neurotherapeutics, 15(9), 1031–1039.

Tian, Y., Du, J., Spagna, A., Mackie, M.-A., Gu, X., Dong, Y., … Wang, K. (2016). Venlafaxine treatment reduces the deficit of executive control of attention in patients with major depressive disorder. Scientific reports, 6, 28028.

Tombu, M., Asplund, C. L., Dux, P. E., Godwin, D., Martin, J. W., & Marois, R. (2011). A unified attentional bottleneck in the human brain. Proceedings of the National Academy of Sciences, 108(33), 13426–13431.

Tombu, M., & Jolicœur, P. (2003). A central capacity sharing model of dual-task performance. Journal of Experimental Psychology: Human Perception and Performance, 29(1), 3.

Trautwein, F.-M., Singer, T., & Kanske, P. (2016). Stimulus-Driven Reorienting Impairs Executive Control of Attention: Evidence for a Common Bottleneck in Anterior Insula. Cerebral Cortex, 26(11), 4136–4147.

Wu, T., Chen, C., Spagna, A., Wu, X., Mackie, M. A., Russell-Giller, S., … Hof, P. R. (2019). The functional anatomy of cognitive control: A domain-general brain network for uncertainty processing. Journal of comparative neurology.

Wu, T., Dufford, A. J., Egan, L. J., Mackie, M. A., Chen, C., Yuan, C., … Fan, J. (2017). Hick-Hyman Law is Mediated by the Cognitive Control Network in the Brain. Cereb Cortex, 1–16. doi: 10.1093/cercor/bhx127

Wu, T., Wang, X., Wu, Q., Spagna, A., Yang, J., Yuan, C., … Fan, J. (2019). Anterior insular cortex is a bottleneck of cognitive control. Neuroimage, 195, 490–504. doi: 10.1016/j.neuroimage.2019.02.042

Xuan, B., Mackie, M.-A., Spagna, A., Wu, T., Tian, Y., Hof, P. R., & Fan, J. (2016). The activation of interactive attentional networks. NeuroImage, 129, 308–319. doi: 10.1016/j.neuroimage.2016.01.017

